# Camouflaging Endovascular Stents with an Endothelial Coat Using CD31 Domain 1-mimetic Peptides

**DOI:** 10.1101/2023.05.01.538569

**Authors:** J. Sénémaud, C. Skarbek, R. Song, Z. Pan, I Lefevre, E. Bianchi, Y. Castier, A. Nicoletti, C. Bureau, G. Caligiuri

**Author notes:** equally contributed (J. Sénémaud, C. Skarbek, C. Bureau, G. Caligiuri).

## Abstract

**Background:** Endovascular stents and flow diverters have become widely used for the treatment of vascular diseases; however, their effectiveness is often limited by the deposition and activation of blood platelets and leukocytes. The foreignness of these devices often triggers pathologic local reactions that impede their integration and compromise their efficacy. In this study, we developed a method to camouflage endovascular stents and flow diverters by coating them with a surface that mimics healthy endothelium, in order to promote more effective device integration and prevent the activation of blood cells.

**Methods:** We designed peptides using domain 1 and/or domain 2 of human CD31 and synthesized the chosen peptide to coat clinical-grade nitinol flow diverters and cobalt chromium balloon expandable stents. The coated stents were implanted in adult rabbits and included control groups of uncoated devices and drug-eluting CoCr stents. The rabbits were monitored for 60 days, during which we assessed the integration of the devices under a physiologic confluence of endothelial cells.

**Results:** Our results demonstrated that the stents coated with the CD31-Domain 1 mimicking peptide promoted a smooth integration of the devices under a physiologic confluence of endothelial cells. By day 7, the coated stents were entirely covered by a smooth endothelium, unlike bare-metal and drug-eluting stents which remained largely exposed to the flowing blood. By day 60, the coated stents demonstrated superiority over both bare-metal and drug-eluting stents, as they resulted in the formation of a “neo-arterial” wall at the entrance of the aneurysmal sac.

**Conclusion:** Our method provides a promising step towards the development of more effective and biocompatible endovascular devices. The CD31 domain 1 coating prevented the pathologic local reaction at the site of stent implantation and promoted faster and more effective device integration. Further studies are necessary to investigate the efficacy and safety of CD31 domain 1 coatings on a larger scale, as well as their long-term durability and potential clinical applications.

## Introduction

Endovascular devices, such as stents and flow diverters, can prolong the activation of blood platelets and leukocytes at the site of implantation, which disturbs the healing of the treated arterial segment and leads to chronic inflammation and neointima development. Previous studies have shown that immobilizing a synthetic peptide (P8RI) derived from the membrane-proximal portion of CD31, which is capable of rescuing its coinhibitory functions on activated blood leukocytes and platelets, can reduce platelet and leukocyte activation in vitro and improve the healing of the stented artery in vivo ^*1, 2*^.

Based on these findings, we hypothesized that CD31 peptides derived from the first two membrane-distal trans-homophilic domains of CD31 could exert a beneficial effect independent of the need to blunt activated leukocytes and platelets. CD31 comprises six Ig-like extracellular domains, and the membrane-proximal portion is not accessible in resting cells. However, the first two domains are widely exposed by healthy endothelium, where they mediate the physiological mutual recognition with platelets, leukocytes, and endothelial cells themselves ^*3, 4*^.

Therefore, we proposed camouflaging the stents and flow diverters with coatings that mimic these domains, enabling the devices to be directly perceived as healthy endothelium by cells contacting their surface. By masking the foreignness of the device, the objective was to prevent the pathologic local reaction at the site of stent implantation and promote faster and more effective device integration.

## Methods

To design the peptides, we encompassed domain 1 and/or domain 2 of human CD31 ^5^. Based on functional properties determined by preliminary in vitro screening (Figure 1A), one peptide from domain 1 (Figure 1B) was selected. We then synthesized the chosen peptide and covalently coated it onto clinical-grade nitinol flow diverters and cobalt chromium (CoCr) balloon expandable stents using click chemistry following a three-step dip-coating procedure as previously described ^*1, 2*^.

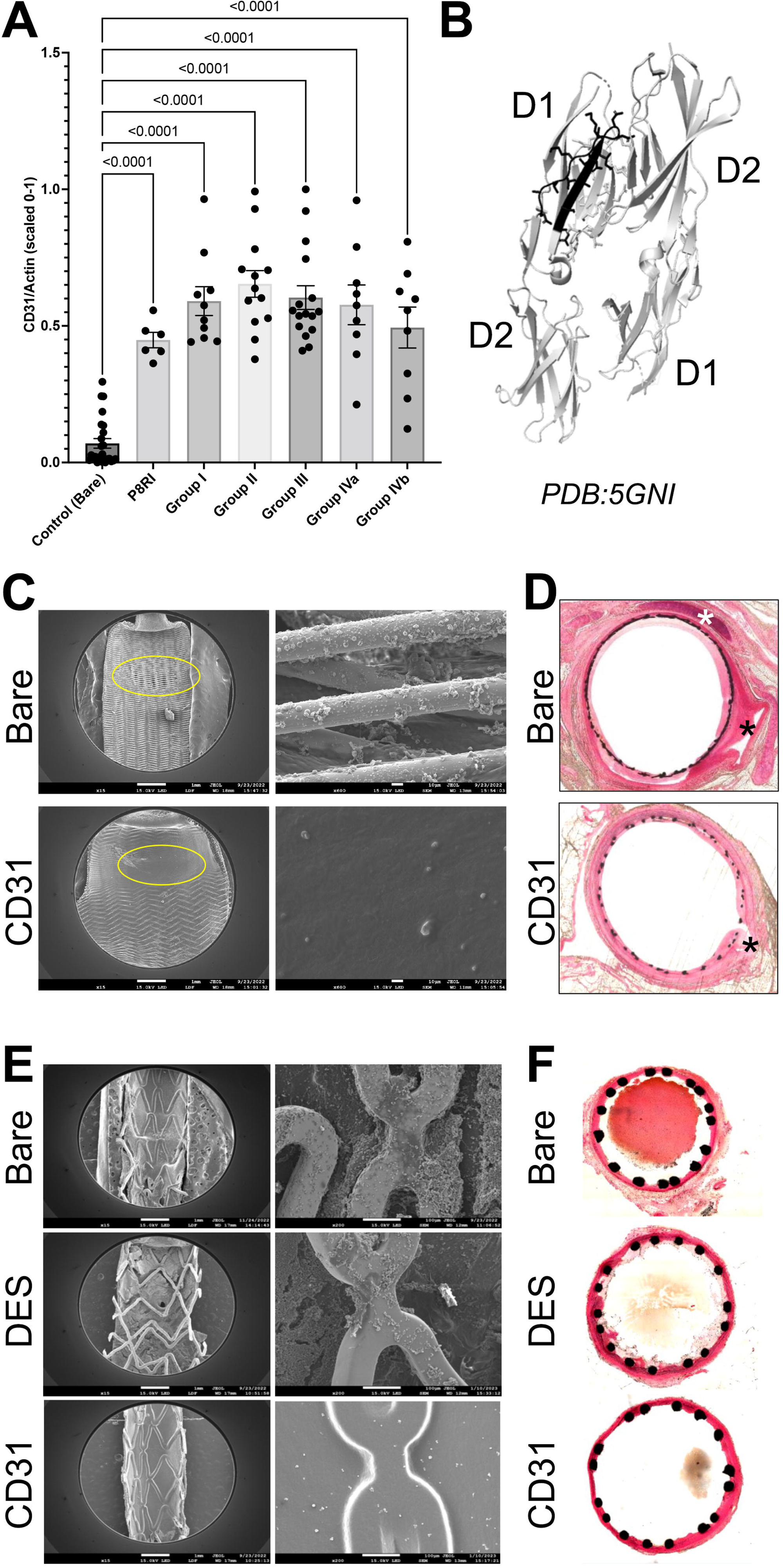
Properties of CD31 Domain 1 endothelial-mimetic peptides. **A**. In vitro testing results of different CD31 peptides spanning trans-homophilic regions on Nitinol flat disks coated with indicated peptides and cultured with human coronary endothelial cells. Group I = aa71-87; Group II = aa107-122; Group III = aa133-158; Group Iva = covalently linked heterodimers of Group II and Group I peptides; Group IVb = covalently linked heterodimers of Group III and Group I peptides. Data are presented as CD31 expression at the cell border to extent of F-actin polymerization (Phalloidin staining), indicating intracellular stress-fibers. CD31-coated surfaces from Group I-III show higher CD31/Actin ratio, indicating a more physiological endothelial cell phenotype compared to bare metal disks. **B**. CD31 Domain 1 peptide used in vivo (black). Illustration of trans-homophilic interaction between CD31 Domains 1 and 2 based on PDB file 5GNI. **C**. SEM image of the luminal face of the right rabbit carotid artery with implanted Bare or CD31 flow diverters 60 days prior. Inset shows higher magnification of region indicated by yellow ellipse. **D**. Representative haematoxylin and eosin-stained resin cross-sections from parallel groups. **E**. SEM image of the luminal face of the rabbit abdominal aorta with implanted Bare, DES, and CD31 CoCr stents 7 days prior. Inset shows higher magnification of stent struts. **F**. Representative haematoxylin and eosin-stained resin cross-sections from parallel groups.

Next, we implanted the experimental stents in adult rabbits, with the flow diverters placed in the subclavian right carotid artery subjected to the elastase saccular aneurysm model, and CoCr stents implanted in the abdominal aorta. We included control groups of uncoated (bare metal) devices and drug-eluting (DES) CoCr stents.

We analyzed the luminal side of experimental arterial segments using scanning electron microscopy (SEM), and resin cross-sections were analyzed using light microscopy.

## Results

In the elastase saccular aneurysm rabbit model, the results showed that flow diverters coated with the CD31-Domain 1 mimicking peptide promoted a smooth integration of the devices under a physiologic confluence of endothelial cells. This resulted in the formation of a “neo-arterial” wall at the entrance of the aneurysmal sac. In contrast, the group implanted with control/bare metal flow diverters exhibited an open inter-strut space at the aneurysmal entry, even two months after implantation, as depicted in Figure 1C. Examination of resin cross-sections revealed that in the control group, fresh blood was still present in the sac (indicated by black asterisk), and flourishing periarterial inflammation was still visible even after two months (indicated by white asterisks). However, in the CD31 group, the arterial tissue appeared to have healed evenly, and the adventitia remained consistently “clean,” as illustrated in Figure 1D.

CoCr stents coated with the selected CD31 peptide demonstrated superiority over both bare-metal and drug-eluting stents. By day 7, CD31-coated CoCr stents were entirely covered by a smooth endothelium, in contrast to drug-eluting and bare-metal stents which remained largely exposed to the flowing blood. Moreover, the endothelium covering the CD31-coated stents remained entirely “clean,” unlike bare-metal stents which often displayed signs of local thrombo-inflammation over the endothelialised struts (refer to Figure 1E). The morphometric analysis of resin cross-sections confirmed the complete absence of thrombo-inflammation on the luminal side of coated stents, as well as the absence of adventitial inflammatory reactions commonly observed with bare metal devices (Figure 1F).

## Conclusions

Our results suggest that the use of peptides that replicate the membrane-distal portion of CD31, which is prominently exposed on the inner side of healthy vessels, can mimic the surface of a healthy endothelium and prevent the deposition and activation of blood platelets and leukocytes that are abnormally prolonged by the presence of the device (foreign body) at sites of endovascular stent and flow diverter implantation. The Domain-1 endothelial-mimetic coating used in this study promotes a rapid and durable integration of the devices beneath a physiologic endothelium. Further studies are necessary to investigate the efficacy and safety of CD31 domain 1 coatings on a larger scale, as well as their long-term durability and potential clinical applications. Overall, this study provides a promising step towards the development of more effective and biocompatible endovascular devices.

